# Mechanical, rheological, and sensory characterization of lion’s mane mushroom steak

**DOI:** 10.64898/2026.01.19.700477

**Authors:** Skyler R. St. Pierre, Lucas Boyle, Thibault Vervenne, Ethan C. Darwin, Marie A. Goodson, Manuel Palomares, Nancy Zhang, Ellen Kuhl

## Abstract

Mushrooms are increasingly recognized as delicious, nutritious, and sustainable foods, with an intrinsic umami flavor and fibrous microstructure that can approximate meat-like texture. Among them, lion’s mane mushroom has emerged as a promising candidate for whole-cut meat alternatives. Yet, its mechanical, rheological, and sensory properties remain largely unquantified. Here we show that a minimally processed lion’s mane mushroom steak exhibits distinctive mechanical, rheological, and sensory characteristics that position it favorably among existing meat alternatives. Despite its pronounced fibrous morphology, lion’s mane steak behaves predominantly as an isotropic material under both mechanical loading and rheological testing, with elastic stiffnesses of *E* = 33.2 kPa and *E* = 34.8 kPa in-plane and cross-plane. A fundamental challenge in alternative protein development is to understand how these measurable physical properties relate to human texture perception. In a complementary sensory survey, *n* = 21 participants ranked lion’s mane steak as more fatty, fibrous, moist, and meaty than eight animal- and plant-based comparison meats. Strikingly, our perceived sensory softness correlates inversely with our experimentally measured mechanical stiffness (τ=-0.60, p = 0.02) and rheological loss modulus (τ=-0.56, p = 0.03). Taken together, our results demonstrate that lion’s mane steak combines favorable mechanical performance with desirable sensory attributes and provide a mechanistic link between physics and taste. Our findings highlight lion’s mane mushroom as a compelling whole-cut alternative protein and underscore the value of integrated mechanical–sensory characterization for rational food design.

## 1. Introduction

Mushrooms are poised to replace meat in people’s diets due to their rich nutritional profile, health-promoting bioactive compounds, umami flavor and fibrous texture, and low environmental footprint [1]. Mushrooms are the fruiting body of the fungus that is visible aboveground, while mycelium are the roots below ground [2]. Mycoprotein, which is made from mycelium, requires more processing to act as a meat substitute than using the fruiting body directly [3]. The lion’s mane mushroom, *Hericium erinaceus*, is widely consumed for its nutritional content [4] and anti-inflammatory, antioxidative, and immunostimulating properties [5], although these health benefits require further human studies to fully validate [6]. While powdered [7] and chopped [8] lion’s mane has been investigated as a potential meat substitute component, the mechanical, rheological, textural, and sensory properties of whole lion’s mane are not well studied.

In this study, we evaluate the mechanical, rheological, textural, and sensory characteristics of lion’s mane mushroom steak. We perform tension, compression, and shear tests [9, 10, 11], double-compression texture profile analysis [12], and oscillatory rheology [13] for both in-plane and cross-plane directions.

We also conducted a sensory survey with twelve questions focused on the mechanical features of meat with twenty-one participants [14]. We compare the performance of the lion’s mane steak to eight commercially available meat products [14]. Finally, using statistical analysis, we evaluate the ability of mechanics, rheology, and texture profile analysis metrics to predict sensory perception. Overall, we seek to answer the question: *Can mushroom steak mimic the mechanical and sensory features of processed animal- and plant-based meat products?*

## 2. Methods

In this study we test the *fungi-based steak* Lion’s Mane Mushroom Steak (OMNI, New York, NY) in two orthogonal directions, and assume that the *in-plane direction* of the steak is parallel to the fiber direction, and the *cross-plane direction* of the steak perpendicular to the fiber direction. The product contains, in descending order by weight, water, lion’s mane mushroom, onion, garlic, psyllium seed husk, sunflower oil, salt, yeast extract, black pepper and beet powder. From the nutrition label, one serving of half the package weighs 100 g and contains 1 g or 1% of the daily value for total fat, 18 g or 7% of daily total carbohydrates, with 15 g or 54% of daily dietary fiber, 1 g of protein, and 270 mg or 12% of daily sodium.

We compare OMNI Lion’s Mane Mushroom Steak against three *animal-based meats*, Turkey Polska Kielbasa Sausage (Hillshire Farm, New London, WI), Spam Oven-Roasted Turkey (Spam, Hormel Foods Co, Austin, MN), and Classic Uncured Wieners (Oscar Mayer, Kraft Heinz Co, Chicago, IL), and five *plant-based meats*, Ham-Style Roast Tofurky (Tofurky, Hood River, OR) Vegan Frankfurter Sausage (Field Roast, Seattle, WA), Plant-Based Signature Stadium Hot Dog (Field Roast, Seattle, WA), Organic Tofu Extra Firm (House Foods, Garden Grove, CA), and Organic Firm Tofu (365 by Whole Foods, Austin, TX). We select these eight comparison products from our previous texture profile analysis and rheological tests [13], and our mechanical tests and sensory surveys [14].

### 2.1. Sample preparation

We biopsy punch a total of *n* = 90 cylindrical samples of 8 mm diameter and 10 mm height, *n* = 45 in-plane, parallel to the fiber direction, and *n* = 45 cross-plane, perpendicular to the fiber direction. We also cut a total of *n* = 20 rectangular samples of 10 mm length, 10 mm width, and 20 mm height, *n* = 10 in-plane and *n* = 10 cross-plane. Figure 1 illustrates the sample preparation process.

**Figure 1:**
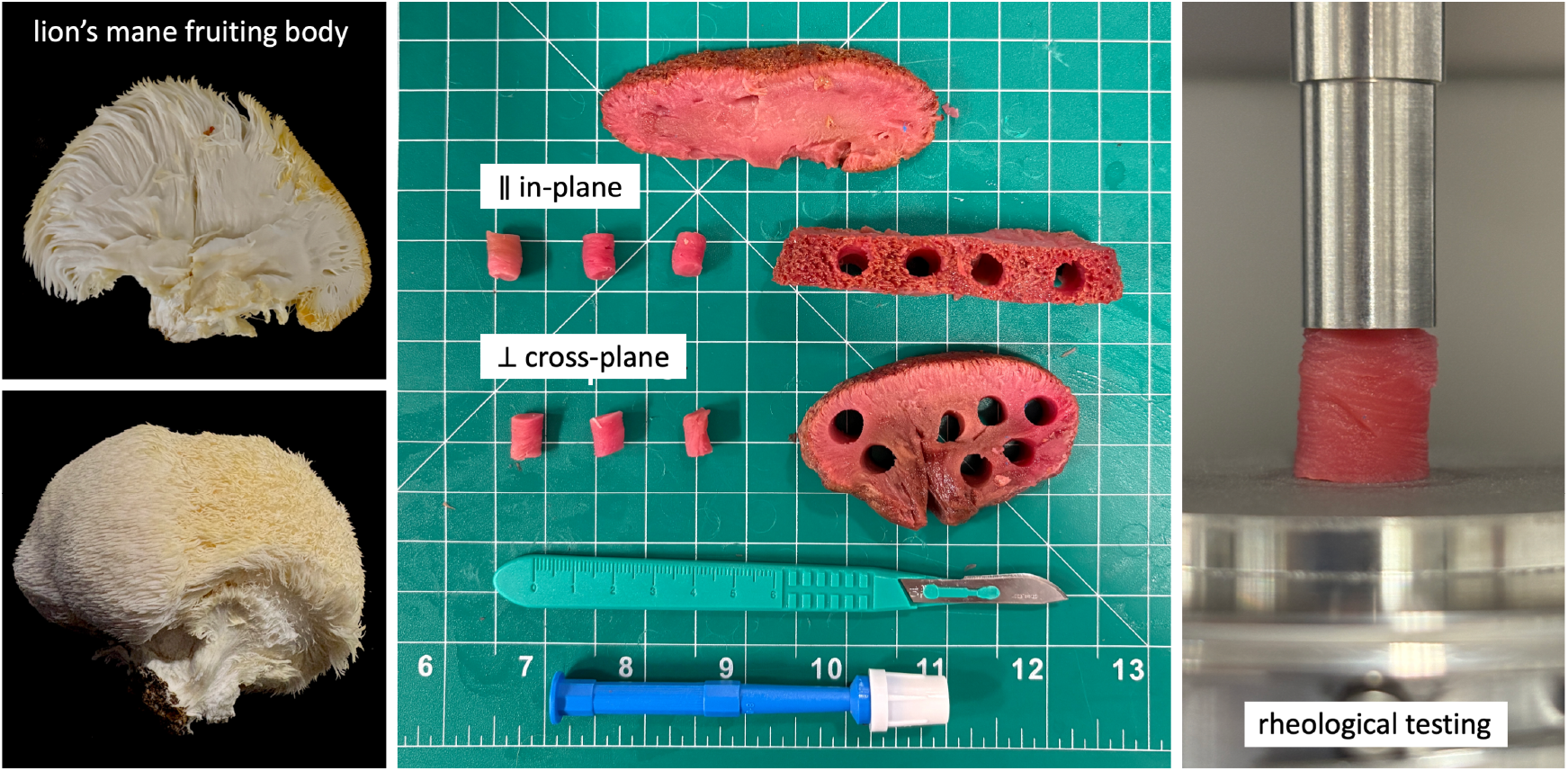
Sample preparation. We test a commercially available fungi-based steak created from the fruiting body of the lion’s mane mushroom in tension, compression, and shear. We use a biopsy punch to extract cylindrical samples with 8 mm diameter and 10 mm height, both in-plane and cross-plane for compression and shear. We cut additional rectangular samples with 10 mm length, 10 mm width, and 20 mm height, in-plane and cross-plane for tension. We test a total of *n* = 110 samples, *n* = 55 in each direction: *n* = 30 quasi-statically in tension, compression, and shear, *n* = 10 in uniaxial double compression at high strain rate, and *n* = 15 in oscillatory shear.

### 2.2. Sample testing

For each direction, in-plane and cross-plane, we prepare *n* = 30 samples for quasi-static tension, compression, and shear testing with ten samples each, *n* = 10 samples for uniaxial double compression testing, and *n* = 15 samples for oscillatory shear testing with five samples for amplitude sweeps and ten for frequency sweeps. We perform all compression and shear tests using an HR20 discovery hybrid rheometer (TA Instruments, New Castle, DE) and mount the samples between a 40 mm diameter base plate and a 8 mm diameter parallel plate, both sand-blasted to avoid slippage. Figure 1 displays our compression and shear test setup with the sample mounted in the rheometer. We apply a consistent mounting force of 0.05 N before testing to ensure proper contact. We use an Instron 5848 (Instron, Canton, MA) for tension testing and glue the samples to glass slides, which we mount in custom testing grips [14].

#### 2.2.1. Tension, compression and shear testing for mechanical analysis

For the quasi-static tension tests, we mount the samples and elongate them to a stretch of λ = 1.1 at a stretch rate of 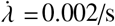, which translates to a total loading time of *t* = 50 s. For the quasi-static compression tests, we mount the samples and compress them to a stretch of λ = 0.8 at a stretch rate of 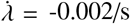, which translates to a total loading time of *t* = 100 s. For the quasi-static shear tests, we mount the samples, apply compression to λ = 0.9, and then shear them to a shear strain of γ = 0.1 at a shear rate of 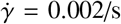, which translates to a total loading time of *t* = 50 s. Figure 2, top left, top right, and bottom left, illustrate the tension stress-stretch, compression stress-stretch, and shear stress-strain behavior where the curves and the shaded regions represent the mean and standard error of the mean across *n* = 10 tests in the in-plane direction parallel to the fibers in light green and in the cross-plane direction perpendicular to the fibers in dark green. The quasi-static tension, compression, and shear tests serve as the basis for the linear mechanical analysis in Section 2.3.

**Figure 2:**
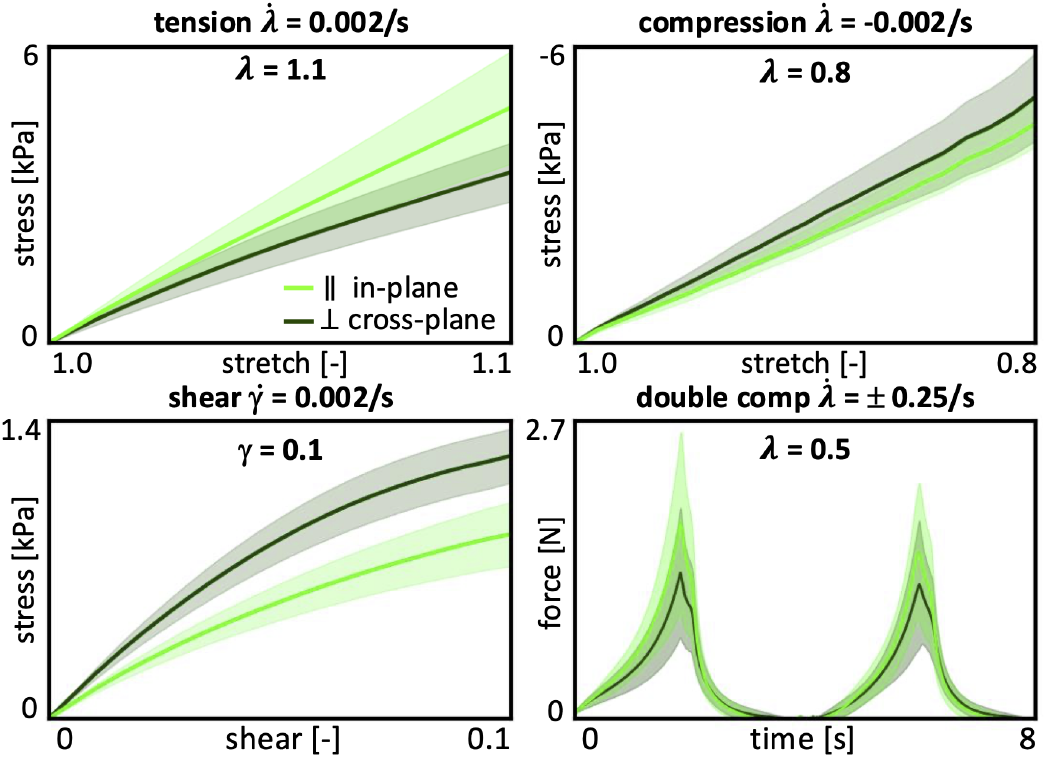
Tension, compression, shear, and double compression testing. Quasi-static tension stress-strain data for 10% strain 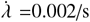 for 50 s, quasi-static compression stress-stretch data for 20% strain at 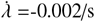 for 100 s, quasi-static shear stress-strain data for 10% shear strain at 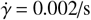 for 50 s, and double compression force-time data for 50% strain at 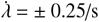 for 8 s. Curves and shaded regions represent the mean and standard error of the mean for quasi-static tests and mean and standard deviation for double compression across *n* = 10 tests in the in-plane and cross-plane directions in light green in dark green.

#### 2.2.2. Double compression testing for texture profile analysis

For the double compression tests, we mount the samples, compress them to λ = 0.5 at a rate of 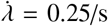, unload the samples at the same rate, and repeat this loading and unloading process a second time, resulting in a total test time of 8 s. Figure 2, bottom right, illustrates the double compression force-time behavior, where the curves and the shaded regions represent the mean and standard deviation of *n* = 10 tests in the in-plane direction parallel to the fibers in light green and in the cross-plane direction perpendicular to the fibers in dark green. The double compression tests serve as the basis for the texture profile analysis in Section 2.4.

#### 2.2.3. Oscillatory shear testing for rheological analysis

For the oscillatory shear tests, we mount the samples and apply a pre-load by compressing them to *λ* = 0.9. This pre-load minimizes slip during oscillatory shear testing and ensures sufficient frictional contact to reduce interfacial motion. First, we perform *n* = 5 amplitude sweep tests, oscillating at 0.5 Hz, with shear amplitudes increasing logarithmically from γ = 0.0001 to *γ* = 0.6. Figure 3, top row, shows the recordings of the amplitude sweep, from which we select the shear amplitude of *γ* = 0.02 to define the linear viscoelastic regime for the subsequent frequency sweeps. We then perform *n* = 10 frequency sweep tests, at an amplitude of *γ* = 0.02, with angular frequencies increasing logarithmically from ω = 0.1 rad/s to ω = 100 rad/s. Figure 3, bottom row, shows the recordings of the frequency sweep, from which we extract the storage modulus *G*^′^, loss modulus *G*^′′^, complex shear modulus *G*^*^, and phase angle *δ*, from the data below an angular frequency of ω < 5 rad/s.We do not observe evidence of systematic slip in the recorded signals of either of the tests. The oscillatory shear tests serve as the basis for the rheological analysis in Section 2.5.

**Figure 3:**
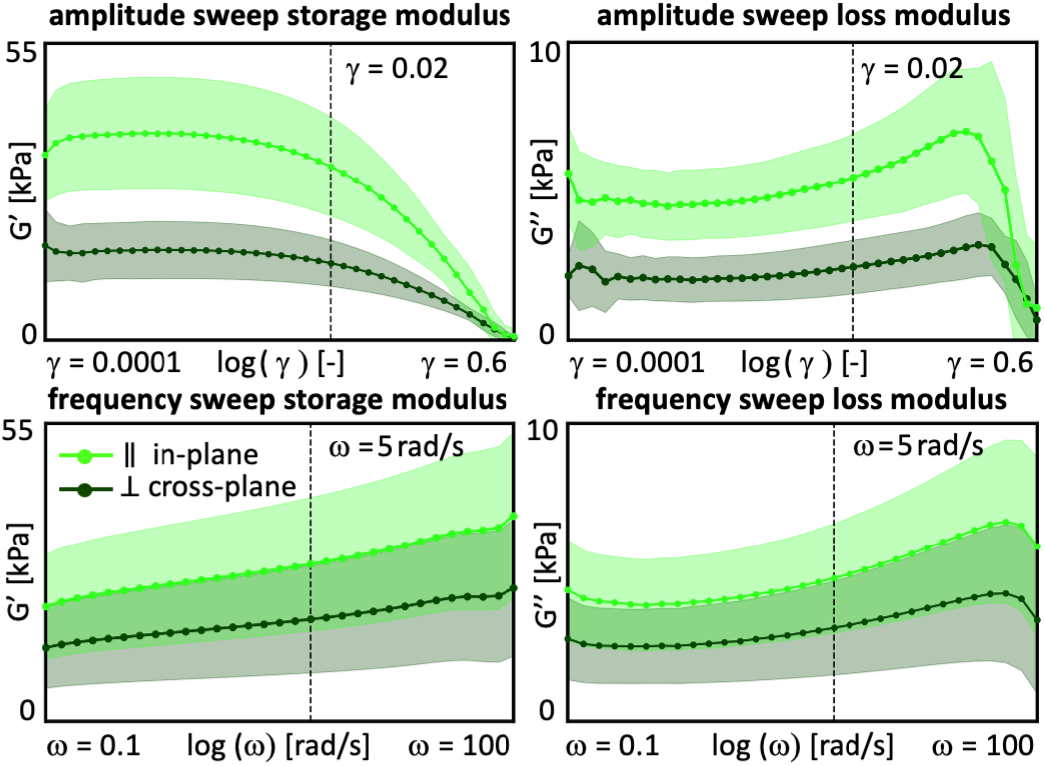
Rheological analysis. We perform amplitude sweep tests, top row, at 0.5 Hz with logarithmically increasing shear amplitudes from *γ* = 0.0001 to *γ* = 0.6. We select the amplitude of *γ* = 0.02 to define the linear viscoelastic regime for the subsequent frequency sweeps. We conduct frequency sweep tests, bottom row, at *γ* = 0.02 with logarithmically increasing angular frequencies from ω = 0.1 rad/s to ω = 100 rad/s. Curves and shaded regions represent the mean and standard deviation of *n* = 5 amplitude sweep and *n* = 10 frequency sweep tests each in the in-plane and cross-plane directions in light green in dark green.

### 2.3. Mechanical analysis

To characterize the *linear elastic* behavior, we perform a linear regression on the quasi-static tension, compression, and shear data throughout 10% tension, 20% compression, and 10% shear to extract the tensile, compressive, shear, and mean stiffnesses [14]. We postulate a linear stress-strain relation, *σ* = *E*. ε, and determine the *tensile* or *compressive* stiffness *E* = (**ε · σ**)/(ε · ε), from the recorded strain-stress pairs {ε; σ }using linear regression. Similarly, we postulate a linear shear stress-strain relation, *τ* = *µ* .γ, convert the shear modulus *µ* into the shear stiffness, *E*_shr_ = 2 [1 + *ν*] *µ* = 3 *µ*, and determine the *shear stiffness*, 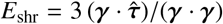, from the recorded shear strain-stress pairs {*γ*; *τ*} . Finally, we average the stiffnesses in tension, compression, and shear to obtain the *mean stiffness, E*_mean_ = (*E*_ten_ + *E*_com_ + *E*_shr_)/3.

### 2.4. Texture profile analysis

To characterize the physics of chewing, we perform a texture profile analysis using the double compression tests from Section 2.2.2. Figure 2, bottom right, illustrates the load profile of the double compression test, which consist of two consecutive cycles of 50% compression. According to standard definitions [15, 16], we denote the peak forces of the first and second loading cycles as *F*_1_ and *F*_2_, the associated loading times as *t*_1_ and *t*_2_, the areas under their loading paths as *A*_1_ and *A*_3_, and the areas under their unloading paths as *A*_2_ and *A*_4_ [13]. We convert the recorded data into stress-strain curves, where the stress σ = *F*/*A* is the recorded force *F* divided by the specimen cross section area *A* = π *r*^2^ = 50.3 mm^2^ with *r* = 4 mm, and the strain 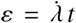 is the applied strain rate multiplied by the time *t* [17]. From the peak force of the first loading cycle *F*_1_, we calculate the peak stress σ_1_ = *F*_1_/*A* at peak strain ε = −50%. From these characteristic values, we extract six texture profile analysis parameters [18]: The *stiffness E* = σ/ε refers to the slope of the stress-strain curve during first compression, when the specimen is compressed to half of its height. The *hardness F*_1_ is associated with the peak force during this first compression cycle. The *cohesiveness* (*A*_3_ + *A*_4_)/(*A*_1_ + *A*_2_) characterizes the material integrity during the second loading and unloading cycle, compared to the first cycle, where a value of one relates to a perfectly intact material, whereas a value of zero relates to complete disintegration. The *springiness t*_2_/*t*_1_ is associated with recovery and viscosity and describes the speed by which the material springs back to its original state after the second cycle compared to the first. The *resilience A*_2_/*A*_1_ is a measure of how well a sample recovers during first unloading path relative to the first loading, where a value of one relates to perfect elasticity whereas a value larger than one indicates plasticity. The *chewiness F*_1_ (*A*_3_ + *A*_4_)/(*A*_1_ + *A*_2_) *t*_2_/*t*_1_, the product of hardness, cohesiveness, and springiness, relates to the resistance of a material during the chewing process, with higher chewiness values indicating that the material is more difficult to chew.

### 2.5. Rheological analysis

To characterize the time-dependent behavior, we perform a rheological analysis using the oscillatory shear tests from Section 2.2.3. From the amplitude and frequency sweeps, we extract four rheological parameters [13]: The *storage modulus G*^′^ = *G*^*^ cos(*δ*) is the apparent or effective stiffness that is a result of both the stiffness of the steak’s solid matrix and the effect of fluid pressure. The *loss modulus G*^′′^ = *G*^*^ sin(*δ*) is the hydraulic dissipation modulus that describes the interstitial fluid flow within the pores of the steak’s solid matrix. The *complex shear modulus G*^*^ = *G*^′^ + *i G*^′′^ is the sum of the storage and loss moduli and describes the effective poroelastic modulus that combines the elastic and dissipative responses of solid and fluid. The *phase angle δ* = tan^−1^(*G*^′′^/*G*^′^) defines the time lag between stress and strain due to delayed fluid distribution and varies between 0^°^≤ *δ* ≤90^°^. The extreme cases of 0^°^ and 90^°^ represent a purely elastic solid that deforms entirely reversibly with all energy stored in the solid matrix and a purely dissipative fluid that deforms entirely irreversibly with all energy lost to fluid flow.

### 2.6. Sensory survey

For the sensory survey, we pan fry the lion’s mane mushroom steaks following the manufacturer’s directions using only a small amount of oil on the pan and then prepare bite-sized samples. We serve the samples immediately after preparation. In accordance with our previous studies, we recruit *n* = 21 participants to participate in the Food Texture Survey. We note that this sample size is relatively small. However, the study design prioritizes consistency with our prior data to enable direct cross-study comparisons [12, 13, 14, 19]. We instruct all participants to eat the samples and rank twelve mushroom steak texture features on a 5-point Likert scale. Each question starts with *this food is* … [20, 21], followed by one of the following features: *soft, hard, brittle, chewy, gummy, viscous, springy, sticky, fibrous, fatty, moist*, and *meaty*. The scale ranges from 5 for strongly agree to 1 for strongly disagree. The participants first eat and rank the steak samples in the cross-plane direction, with the fiber direction perpendicular to the initial chewing direction, and then in the in-plane direction, with the fiber direction parallel to the initial chewing direction. Between the samples, we ask participants to cleanse their palate with water to minimize residual flavor, neutralize their taste, and standardize the sensory environment. We follow our established protocol and do not include randomization, cross-over design, or sensory blinding to enable consistent comparisons with our previous studies [12, 13, 14, 19]. This research was reviewed and approved by the Institutional Review Board at Stanford University under the protocol IRB-75418.

### 2.7. Statistical analysis

We export the raw tension, compression, and shear data and analyze the results in Python 3.9. To quantify to which extent the in-plane and cross-plane features differ from each other, we perform a Welch’s t-test for all texture profile analysis, rheological, and sensory features except for the texture profile analysis features of cohesiveness, springiness and resilience, where we use a Mann-Whitney U test for non-continuous data. We report all possible correlations between rankings of mechanics, texture, rheology, and sensory attributes using Kendall’s rank correlation coefficient, τ. We also report effect sizes using Cohen’s *d* to quantify the magnitude of differences in sensory attributes. In addition, we perform a principal component analysis on the rank-based feature matrix to identify dominant modes of variation and clustering across products in a reduced-dimensional space.

## 3. Results

Table 1 provides the detailed information of the product, the Lion’s Mane Mushroom Steak, its brand, ingredients, and nutrition facts. The table also summarizes the stiffness features, texture profile analysis features, rheological features, and sensory features. All features are reported in the in-plane and cross-plane directions, with their means and standard deviations. In the following figures, we compare all features of the fungi-based steak in-plane (FS^||^) and cross-plane (FS^⊥^) against three animal-based products, animal turkey (AT), animal sausage (AS), and animal hotdog (AH), and against five plant-based products, plant turkey (PT), plant sausage (PS), plant hotdog (PH), extrafirm tofu (ET), and firm tofu (FT) [14].

**Table 1:** Lion’s Mane Mushroom Steak. Product; brand; ingredients; nutrition facts; stiffness; texture profile analysis features; rheological features; sensory features. All features are reported in in-plane and cross-plane directions with means and standard deviations

|  | in-plane \|\| | cross-plane ⊥ |
| --- | --- | --- |
| product | Lion's Mane Mushroom Steak |  |
| brand | OMNI, New York, NY |  |
| ingre-<br>dients | water, lion's mane mushroom, onion, garlic, psyllium seed husk, sunflower oil, salt, yeast extract, black pepper, beet powder |  |
| nutrition<br>facts | serving size of half package contains 1 g total fat, 18 g carbohydrates, 15 g dietary fiber, 1 g protein, and 270 mg sodium |  |
| stiffness | $E_{ten} = 48.9 \pm 35.3$ kPa<br>$E_{com} = 21.5 \pm 7.3$ kPa<br>$E_{shr} = 29.2 \pm 15.1$ kPa<br>$E_{mean} = 33.2$ kPa | $E_{ten} = 36.8 \pm 21.1$ kPa<br>$E_{com} = 24.4 \pm 13.7$ kPa<br>$E_{shr} = 43.2 \pm 15.5$ kPa<br>$E_{mean} = 34.8$ kPa |
| texture<br>profile<br>analysis | stiffness 71.45±33.04 kPa<br>hardness 1.8±0.83 N<br>cohesiveness 0.75±0.11<br>springiness 0.74±0.08<br>resilience 0.45±0.05<br>chewiness 0.99±0.48 N | stiffness 52.68±23.36 kPa<br>hardness 1.33±0.59 N<br>cohesiveness 0.84±0.06<br>springiness 0.77±0.11<br>resilience 0.53±0.09<br>chewiness 0.84±0.35 N |
| rheology | $G' = 25.4 \pm 11.1$ kPa<br>$G'' = 4.2 \pm 1.6$ kPa<br>$G^* = 25.7 \pm 11.2$ kPa<br>$\tan(\delta) = 0.168 \pm 0.022$ | $G' = 16.3 \pm 8.7$ kPa<br>$G'' = 2.7 \pm 1.3$ kPa<br>$G^* = 16.6 \pm 8.8$ kPa<br>$\tan(\delta) = 0.166 \pm 0.014$ |
| sensory<br>survey | soft 3.5±0.9 hard 2.2±1.0<br>brittle 2.0±1.0 chewy 4.0±1.0<br>gummy 3.8±1.0 viscous 2.9±1.3<br>springy 3.3±1.3 sticky 1.9±1.2<br>fibrous 4.4±0.8 fatty 3.4±1.1<br>moist 4.0±0.8 meaty 4.1±1.1 | soft 3.6±1.2 hard 2.4±1.3<br>brittle 1.6±0.8 chewy 4.6±0.5<br>gummy 3.7±1.1 viscous 3.0±1.3<br>springy 4.2±0.9 sticky 2.3±1.2<br>fibrous 4.2±0.9 fatty 3.7±1.1<br>moist 4.1±1.0 meaty 3.9±0.9 |

### 3.1. Mechanical analysis

Figure 4 summarizes the results of the linear regression to characterize the *linear elastic* behavior of the lion’s mane steak in the in-plane and cross-plane directions for quasi-static tension up to 1.1 stretch, compression up to 0.8 stretch, and shear up to 0.1 shear strain. The four bar graphs illustrate the in-plane and cross-plane tensile, compressive, shear, and mean stiffness of the fungi-based steak. All stiffness values are fairly similar, with the stiffnesses in the in-plane direction, E_ten_ = 48.9 ±35.3 kPa, E_com_ = 21.5± 7.3 kPa, E_shr_ = 29.2 ±15.1 kPa, and E_mean_ = 33.2 kPa not significantly different than in the cross-plane direction, E_ten_ = 36.8 ± 21.1 kPa, E_com_ = 24.4 ± 13.7 kPa, E_shr_= 43.2± 15.5 kPa, and E_mean_ = 34.8 kPa. The absence of directional dependence suggests that the underlying microstructural hyphal network lacks a dominant structural alignment. For comparison, we also include the values of three animal-based products, animal turkey, animal sausage, and animal hotdog, and five plant-based products, plant turkey, plant sausage, plant hotdog, extrafirm tofu and firm tofu [14]. The mean stiffness of lion’s mane steak is approximately 20% the stiffness of plant turkey at 223.7 kPa, 50% the stiffness of plant sausage and animal turkey at 103.9 kPa and 71.1 kPa, about equal to plant hotdog, animal sausage, and animal hotdog at 38.2 kPa, 37.5 kPa, and 37.1 kPa, and only slightly stiffer than firm tofu and extra firm tofu at 26.3 kPa and 24.3 kPa.

**Figure 4:**
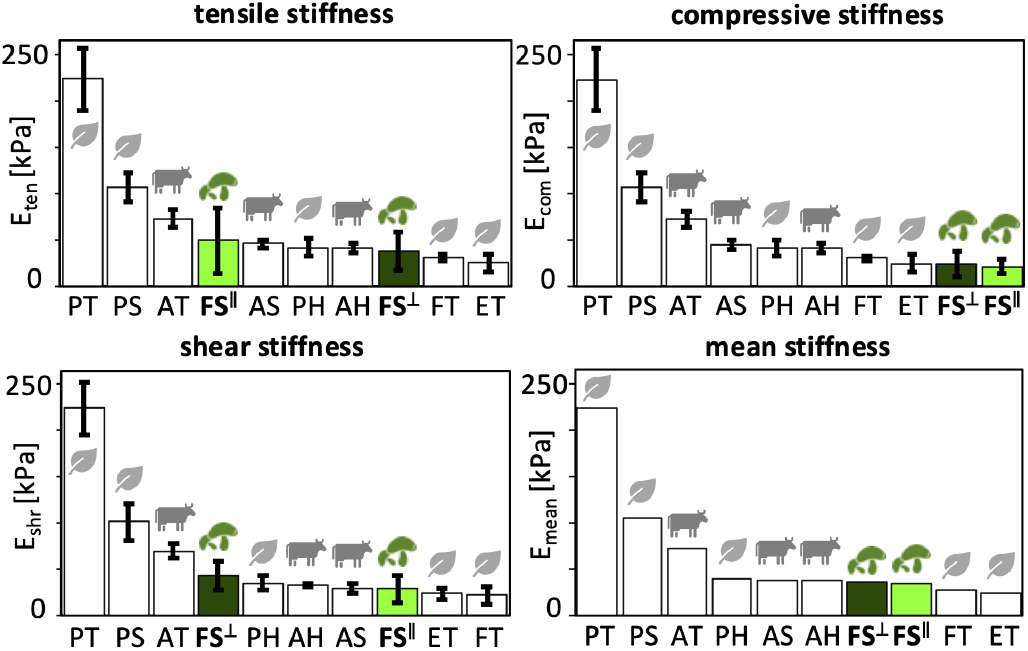
Mechanical analysis. Tensile stiffness, compressive stiffness, shear stiffness, and mean stiffness. In-plane and cross-plane values for OMNI lion’s mane fungi-based steak, highlighted in light and dark green, compared to animal- and plant-based products previously tested [14]. FS^||^ fungi steak in-plane, FS^⊥^ fungi steak cross-plane, AT animal turkey, AS animal sausage, AH animal hotdog, PT plant turkey, PS plant sausage, PH plant hotdog, ET extrafirm tofu, and FT firm tofu.

### 3.2. Texture profile analysis

Figure 5 summarizes the results of texture profile analysis of the lion’s mane steak for the in-plane and cross-plane directions using a double compression to 50% at a strain rate of 25%/s. The six bar graphs illustrate the in-plane and cross-plane stiffness, hardness, cohesiveness, springiness, resilience, and chewiness of the fungi-based steak: The *stiffness* is the slope of the stress-strain curve during the first compression; the *hardness* is the peak force during this first compression cycle; the *cohesiveness* is the material integrity during the second cycle, compared to the first cycle; the *springiness* is the relative time by which the sample recovers; the *resilience* is a measure of how well the sample recovers during the first unloading path relative to the first loading path; and the *chewiness* is the product of hardness, cohesiveness, and springiness. Notably, for all six features, the cross-plane values and in-plane values are not statistically significantly different. Strikingly, stiffness, hardness, resilience, and chewiness of the lion’s mane steak are all lower than those of the animal-based meats, while cohesiveness and springiness rank in the middle of the animal- and plant-based products [13].

**Figure 5:**
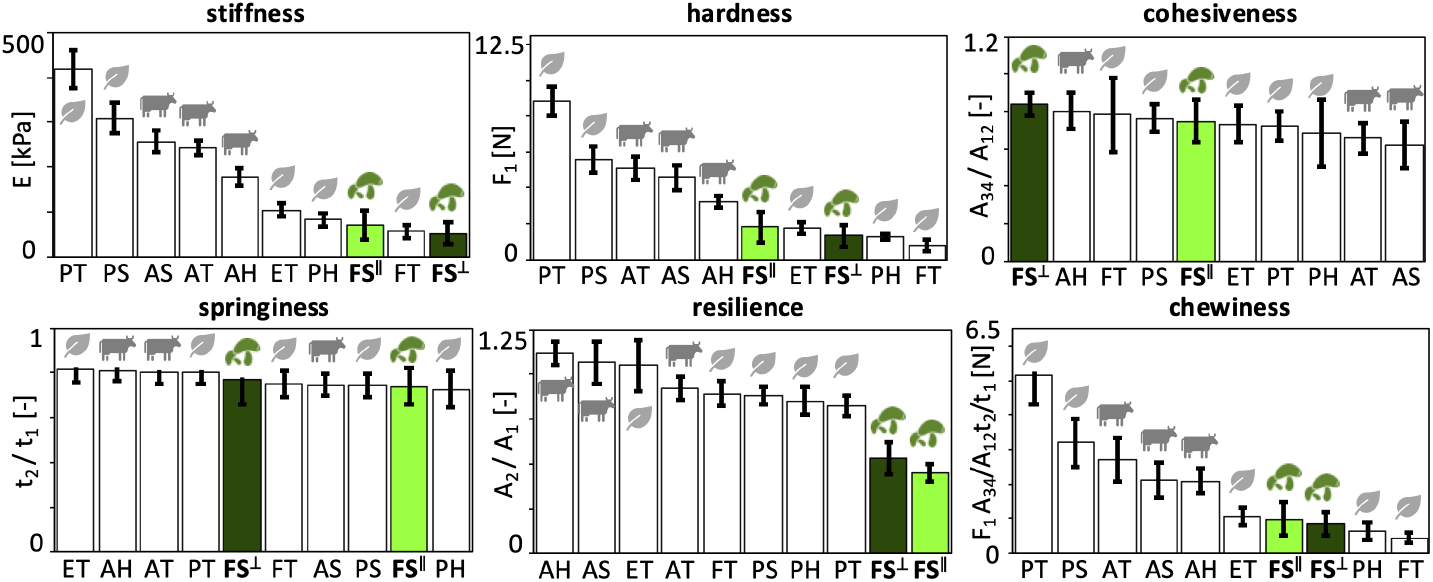
Texture profile analysis. Stiffness, hardness, cohesiveness, springiness, resilience, and chewiness. In-plane and cross-plane values at a loading rate 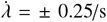, for a total time of 8 s, for lion’s mane fungi-based steak, highlighted in light and dark green, compared to animal- and plant-based products. FS^||^ fungi steak in-plane, FS^⊥^ fungi steak cross-plane, AT animal turkey, AS animal sausage, AH animal hotdog, PT plant turkey, PS plant sausage, PH plant hotdog, ET extrafirm tofu, and FT firm tofu. In-plane and cross-plane values are not statistically different for any of the texture profile analysis parameters.

### 3.3. Rheological analysis

Figure 6 summarizes the rheological analysis of the lion’s mane steak for the in-plane and cross-plane directions. The four bar graphs illustrate the in-plane and cross-plane storage moduli, loss moduli, complex shear moduli, and phase angles, of lion’s mane steak compared to the animal- and plant-based products [13]. We recorded in-plane and cross-plane *storage moduli* of *G*^′^ = 25.4± 11.1 kPa and *G*^′^ = 16.3± 8.7 kPa, *loss moduli* of *G*^′′^ = 4.2 ±1.6 kPa and *G*^′′^ = 2.7± 1.3 kPa, *complex shear moduli* of *G*^*^ = 25.7± 11.2 kPa and *G*^*^ = 16.6± 8.8 kPa, and *phase angles* of tan(*δ*) = 0.168 ±0.022 and tan(*δ*) = 0.166± 0.014. Notably, only the loss moduli are significantly different in-plane and cross-plane with *p* < 0.05. The storage moduli of *G*^′^ = 25.4 kPa and 16.3 kPa quantify the elastic or recoverable part of the mechanical response and suggest that fungi-based steak is stiffer and more elastic in-plane than cross-plane. The loss moduli of *G*^′′^ = 4.2 kPa and 2.7 kPa, quantify the hydraulic or dissipative part of the mechanical response and suggest that fungi-based steak dissipates more energy in-plane than cross-plane, although both values are relatively low compared to the storage moduli *G*^′^ . The phase angles of tan(*δ*) = 0.168 and 0.166, approximately 9.5^°^, measure the lag between stress and strain and suggest a very weak damping and strong elasticity both in-plane and cross-plane. Interestingly, the storage, loss, and complex shear moduli of lion’s mane steak all lie well within the range of the animal- and plant-based products. At the same time, the phase angles are notably lower, suggesting a more elastic response.

**Figure 6:**
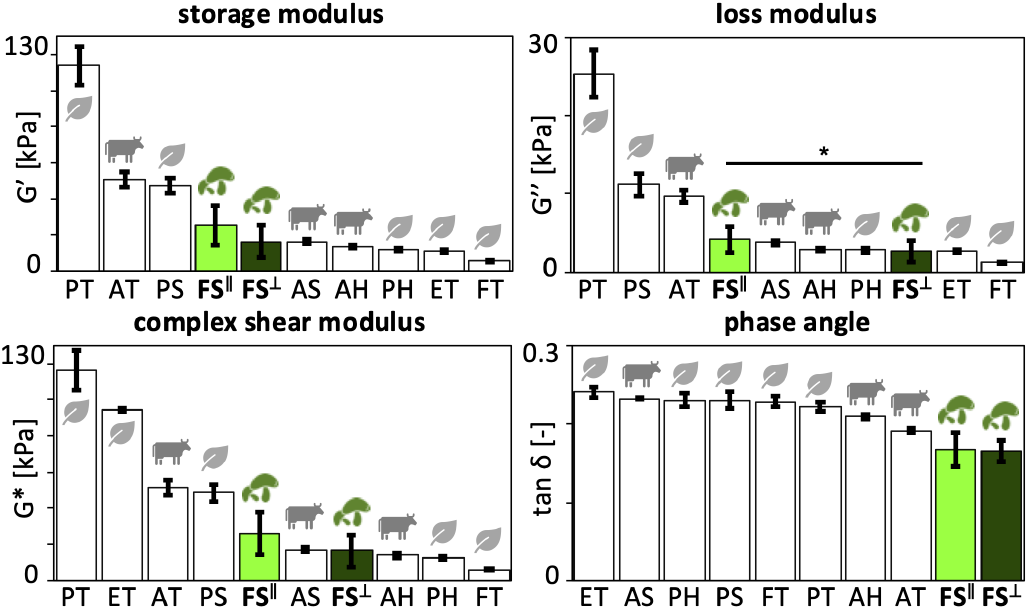
Rheological analysis. Storage modulus, loss modulus, complex shear modulus, and phase angle. In-plane and cross-plane values for lion’s mane steak, highlighted in light and dark green, compared to animal- and plant-based products. FS^||^ fungi steak in-plane, FS^⊥^ fungi steak cross-plane, AT animal turkey, AS animal sausage, AH animal hotdog, PT plant turkey, PS plant sausage, PH plant hotdog, ET extrafirm tofu, and FT firm tofu. Statistical significance between in-plane and cross-plane is indicated as * for *p* ≤ 0.05.

### 3.4. Sensory survey

Figure 7 summarizes the sensory survey results of lion’s mane steak for the in-plane and cross-plane directions. The twelve bar graphs illustrate the in-plane and cross-plane softness, hardness, brittleness, chewiness, gumminess, viscosity, springiness, stickiness, fibrousness, fattiness, moistness, and meatiness of the lion’s mane steak compared to the animal- and plant-based products [14]. Both directions of the lion’s mane steak achieve comparable responses across most attributes, with no significant differences except for *chewiness* and *springiness*, where the cross-plane direction ranks higher (*p* < 0.05). For *softness, hardness*, and *brittleness*, lion’s mane steak ranks well within the animal- and plant-based products. For *springiness* and *stickiness*, the cross-plane direction is the highest ranked, while in-plane is ranked in the middle. Strikingly, across all other attributes, *chewiness, gumminess, viscosity, fibrousness, fattiness, moistness*, and *meatiness*, both directions of lion’s mane steak score higher than all eight comparison products:

**Figure 7:**
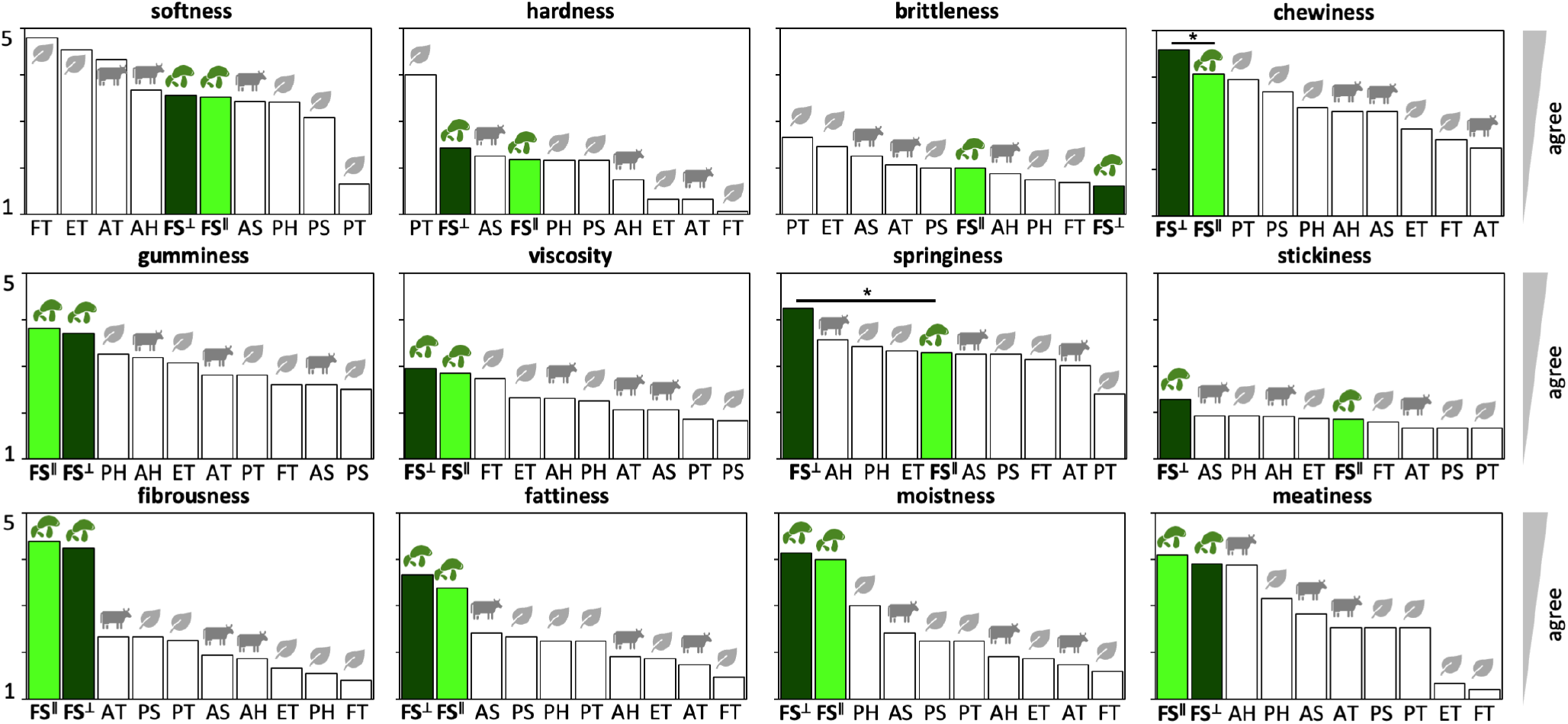
Sensory survey. Softness, hardness, brittleness, chewiness, gumminess, viscosity, springiness, stickiness, fibrousness, fattiness, moistness, and meatiness. In-plane and cross-plane values for lion’s mane steak, highlighted in light and dark green, compared to animal- and plant-based products. FS^||^ fungi steak in-plane, FS^⊥^ fungi steak cross-plane, AT animal turkey, AS animal sausage, AH animal hotdog, PT plant turkey, PS plant sausage, PH plant hotdog, ET extrafirm tofu, and FT firm tofu. Statistical significance between in-plane and cross-plane is indicated as * for *p* ≤ 0.05.

For *fibrousness* and *moistness*, both directions exceed all other products (*p* < 0.001) and form a clearly separated top tier with large effect sizes (*d*≈ 1.0–3.6). For *fattiness*, the same trend holds with reduced separation from the next-best product, animal sausage (*p* < 0.05). For *meatiness*, lion’s mane steak remains among the top performers, but becomes statistically indistinguishable from animal hotdog (*d* < 0.3), indicating a broader high-performing cluster. Ranking robustness reveals strong agreement across *fibrousness, fattiness*, and *moistness* (Kendall’s *τ* ≈0.8–0.95), consistent with a shared structural–sensory axis, while *meatiness* shows reduced concordance (*τ* ≈0.6–0.7). Despite the moderate sample size of *n* = 21, the within-subject design and large effect sizes (*d* ≈ 1.0–3.6) provide sufficient statistical power to support the reported differences.

### 3.5. Correlations between all 26 features

Figure 8 summarizes all possible Kendall’s τ rank correlations between the mechanics, texture profile analysis, rheological analysis, and sensory perception data. The rankings of the four mechanical features, tensile, compressive, shear, and mean stiffness, are from Figure 4; the texture features, stiff, hard, cohesive, springy, resilient, and chewy, are from Figure 5; the rheological features, storage modulus, loss modulus, complex modulus, and phase angle, are from Figure 6; the sensory features, soft, hard, brittle, chewy, gummy, viscous, springy, sticky, fibrous, fatty, moist, and meaty, are from Figure 7. The left heatmap shows the values of Kendall’s *τ*, with dark red, *τ* = +1.0, indicating a perfect positive correlation and dark blue, *τ* = −1.0, a perfect negative correlation. The right heatmap shows the corresponding P-value for each correlation, with dark red indicating no statistical significance, *p* = 1, and dark blue indicating statistically significant, *p* = 0. Both heatmaps are symmetric along the diagonal. Unsurprisingly, there are several strong correlations within each of the four feature groups, mechanics, texture, rheology, and sensory perception, in the four large square blocks around the diagonal, confirming, for example, that the tensile, compressive, shear, and mean stiffnesses are not independent from one another. Similarly, we observe correlations between the three physics-based feature groups, mechanics, texture, and rheology, for example confirming that the *mean stiffness* from mechanics is positively correlated with the *stiffness* from texture profile analysis with Kendall’s rank correlation coefficient of τ = 0.60 (*p* = 0.02). Yet, we are most interested in correlations between the three physical features and sensory perception, in the right and bottom blocks: First, and probably most remark-able, we find that the *softness* from sensory perception is inversely correlated with the *mean stiffness* from mechanics with Kendall’s rank correlation coefficient of *τ* = −0.60 (*p* = 0.02), while the sensory *hardness* is not significantly correlated with any mechanical features. Second, the *brittleness* from sensory perception is positively correlated with the *stiffness* from texture profile analysis with Kendall’s rank correlation coefficient of *τ* = 0.64 (*p* = 0.01). Third, the *softness* from sensory perception is negatively correlated with the *loss modulus* from rheology with Kendall’s rank correlation coefficient of *τ* = −0.56 (*p* = 0.03).

**Figure 8:**
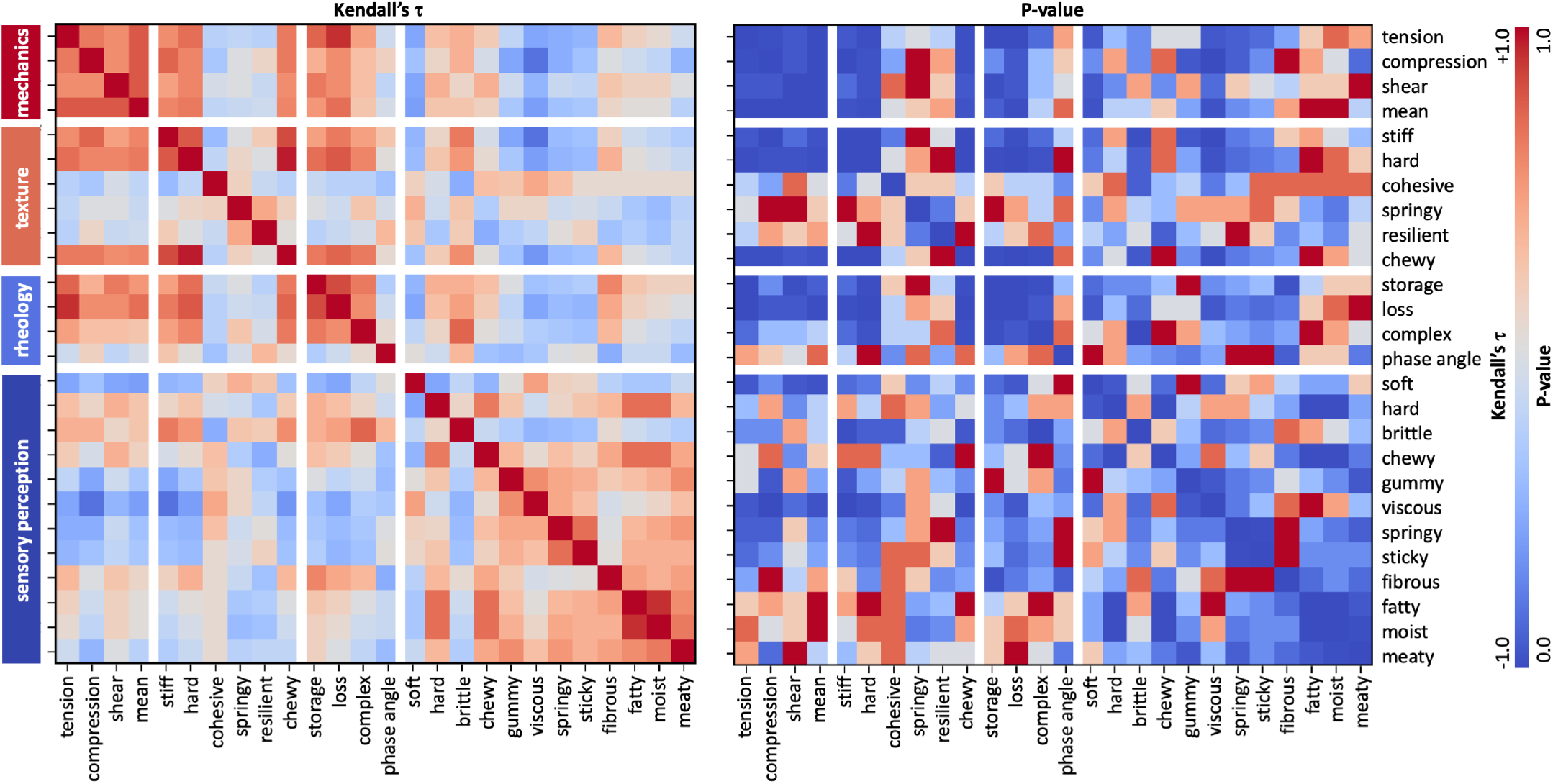
Kendall’s τ rank correlations between mechanics, texture profile analysis, rheology, and sensory perception. Kendall’s rank correlation coefficient, τ, is computed for the ordered results from Figures 4, 5, 6, and 7 of lion’s mane steak and animal- and plant-based products ranging from dark red, τ = +1.0, indicating perfect positive correlation to dark blue, τ= − 1.0, perfect negative correlation. The right heatmap shows the corresponding p-values ranging from dark red, *p* = 1.0, no statistical significance, to dark blue, *p* = 0.0, statistically significant.

### 3.6. Principal component analysis

Figure 9 highlights the results of the principal component analysis of the rank-based product profiles. The first two principal components explain 23.7% and 19.2% of the variance. The first principal component, PC1, aligns with mechanical stiffness, textural hardness and chewiness, and rheological loss and complex moduli. As such, it separates stiffer, highly structured products along the positive direction from softer, more compliant products along the negative direction. The second principal component, PC2, captures sensory attributes, with strong positive loadings for viscosity, gumminess, moistness, fattiness, fibrousness, and meatiness. Lion’s mane mushroom steak, both in-plane FS^||^ and cross-plane FS^⊥^, clusters in the upper-right quadrant with high PC2 scores, indicating strong sensory-rich attributes combined with moderate mechanical stiffness. In contrast, the animal-based and plant-based products distribute primarily along PC1, with stiffer products located in the lower-right quadrant and softer products shifting toward the left side of the plot.

**Figure 9:**
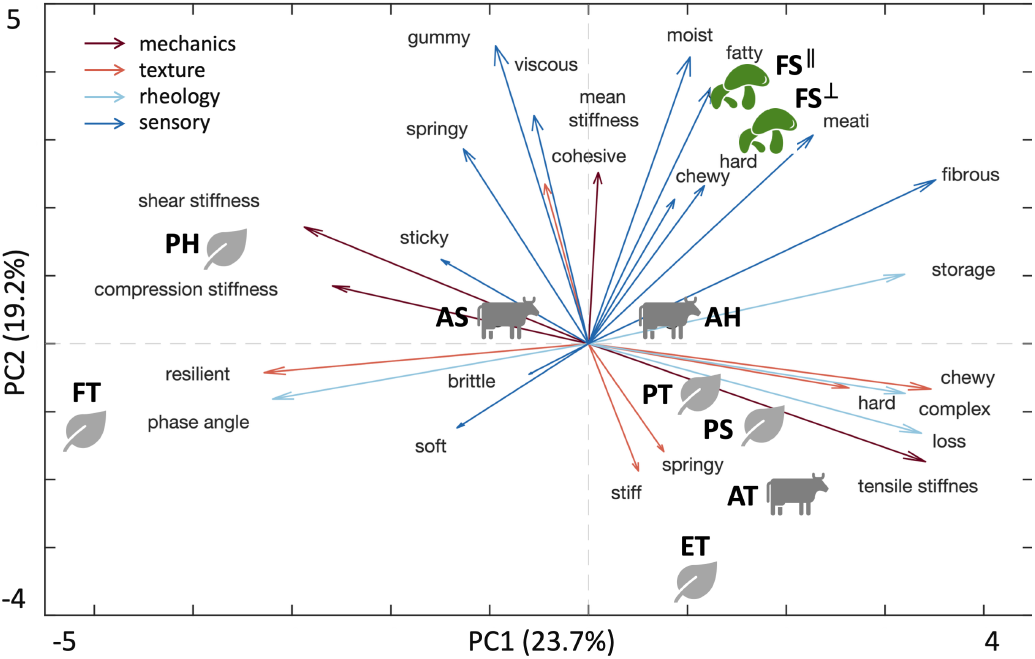
Principal component analysis of rank-based product profiles. The first two principal components explain 23.7% and 19.2% of the variance. Arrows denote feature loadings grouped into mechanics, texture profile analysis, rheology, and sensory attributes. Products cluster by class, with lion’s mane mushroom steak (FS^||^,FS^⊥^) separating along the sensory-richness axis defined by moistness, fattiness, meatiness, and fibrousness, while mechanically stiff products (AT,PS,PT) align along the positive PC1 direction. FS^||^ fungi steak in-plane, FS^⊥^ fungi steak cross-plane, AT animal turkey, AS animal sausage, AH animal hotdog, PT plant turkey, PS plant sausage, PH plant hotdog, ET extrafirm tofu, and FT firm tofu.

## 4. Discussion

We have successfully quantified the mechanical, rheological, and sensory properties of lion’s mane mushroom steak and established direct links between physical measurements and perceived texture. Our results suggest a direct physical basis for several sensory attributes: *Softness* scales inversely with elastic stiffness, consistent with the negative correlation between perceived softness and mean stiffness. *Viscous dissipation* is reflected directly through the loss modulus, where lower dissipation enables faster recovery and a softer texture. *Brittleness* increases with texture profile stiffness, as stiffer materials accumulate higher stresses prior to fracture. These relationships provide a physical basis for sensory perception and show that mechanics and rheology can predict key aspects of texture, consistent with established oral processing models that relate deformation, fracture, and fluid flow to texture perception.

### Microstructure explains weak anisotropy

Recent studies have linked macrostructure, microstructure, and mechanical anisotropy in meat analogues [22]. The lion’s mane steak in this study originates from the fruiting body of *Hericium erinaceus*, which forms a three-dimensional network of branched hyphae that creates a highly porous, water-rich scaffold [23, 27]. Individual hyphae consist of tubular filaments with cell walls composed of chitin, β-glucans, and proteins [24], which define the solid elastic behavior, while the interstitial fluid governs viscoelastic and poroelastic behavior [12]. Unlike mycelium-based materials, which can develop highly aligned fiber architectures [25], the fruiting body organizes into a radially branching, interwoven structure that lacks a dominant macroscopic orientation. This interwoven structure helps explain the weak mechanical anisotropy across our study: The stiffness values across tension, compression, and shear in Figure 4 do not display statistically significant differences between the in-plane and cross-plane directions (p > 0.05), consistent with an effectively isotropic response. The quasi-static tension and compression responses in Figure 2 remain nearly identical in both directions and the texture profile analysis in Figure 5 reveals no directional dependence. This suggests that the branched hyphal network distributes load isotropically, despite its fibrous appearance [26]. Only the loss modulus in Figures 3 and 6 shows a measurable difference between both directions, with higher dissipation in-plane. This is in stark contrast with skeletal muscle, where muscle fibers align within a collagen-rich extracellular matrix [28, 29], and generate a pronounced anisotropy that defines the characteristic fibrous texture of animal steak. Instead, lion’s mane mushroom steak relies on a porous, isotropic hyphal network, which generates comparable mechanical properties in both directions.

### Sensory data reveal that lion’s mane is a promising meat substitute

Figure 7 compares our sensory perception of lion’s mane steak to our prior study on eight different animal- and plant-based meat products with the same survey methods [14]. Participants ate samples with initial chewing directions in-plane and cross-plane. We found that only springiness and chewiness were significantly different between both directions, with cross-plane higher for both features. Strikingly, the lion’s mane steak is ranked as the most fibrous, fatty, moist, and meaty for both in-plane and cross-plane directions compared to all eight comparison products, animal turkey, animal sausage, animal hot-dog, plant turkey, plant sausage, plant hotdog, extrafirm tofu, and firm tofu. The fibrousness of lion’s mane steak more than twice of those of all other meats, which is unsurprising given the minced and chopped nature of the comparison products. However, the high fatty, moist, and meaty scores suggest that lion’s mane mushroom steak is a suitable animal meat substitute.

### Mechanics, texture profile analysis, and rheology features are predictors of sensory perception

Figure 8 reveals the Kendall’s τ rank correlations between mechanics, texture profile analysis, rheology, and sensory perception. Recent studies have used similar correlation matrices to identify correlations between macrostructure, microstructure, and mechanical anisotropy in meat analogues [22]. In our case, only a subset of combinations reaches the level of statistical significance; yet, these feature pairings demonstrate the power of mechanical experiments to predict sensory perception. Notably, our sensory perception of softness is inversely correlated with the mean stiffness from mechanical tests at τ = −0.60 (*p* = 0.02) and with the loss modulus from rheological tests at τ = −0.56 (*p* = 0.03), and our sensory perception of brittleness is positively correlated with the stiffness from texture profile analysis at τ = +0.64 (*p* = 0.01). This highlights the scientific relevance of physics-based testing as a repeatable, unbiased, and objective alternative to sensory surveys with human panels.

### Multivariate structure reveals partially independent axes

While pairwise correlations identify predictive relationships between individual features, principal component analysis captures how these features jointly organize across products. The principal component analysis in Figure 9 identifies two partially independent axes of variation: a mechanics-dominated stiffness axis and a sensory-richness axis. Lion’s mane mushroom steak separates most strongly along the sensory axis, indicating that favorable meat-like perception emerges independently of maximum mechanical stiffness. This decoupling suggests that whole-cut fungi-based foods achieve meat-like perception [1] through microstructural and fluid-mediated mechanisms, rather than through high structural rigidity alone. These findings align with established oral processing models that relate deformation, fracture, and fluid flow to texture perception [20, 21].

### Structured data enable predictive design of fungi-based foods

Quantitative links between physics-based testing and sensory perception establish a foundation for data-driven food design [30]. These relationships support predictive models that reduce reliance on time-consuming sensory surveys and enable inverse design of foods with targeted microstructural and textural properties [31]. The main limitation remains the scarcity of structured datasets that span mechanics, rheology, and perception [32]. As a first step, Figure 8 presents a correlation matrix across 26 food features using Kendall’s τ, a simple and interpretable statistical measure. Open datasets like these expand access to data-driven design tools across academia, industry, and startups [33]. This framework can accelerate the development of foods that integrate sustainability, nutrition, and sensory performance.

## 5. Conclusion

Fungi are a resilient, fast-growing, and nutritious source of food. Here we investigated the potential of lion’s mane mushroom steak as an animal meat substitute with almost no additional processing and minimal added ingredients. The product is made of the fruiting body of lion’s mane, *Hericium erinaceus*, simply sliced into steak-like pieces. Under quasi-static tension, compression, and shear testing, rheological analysis, and high strain rate texture profile analysis, the lion’s mane steak behaves predominantly as an isotropic material with only the loss modulus exhibiting a significant fiber-direction dependence. Physics-based testing puts lion’s mane steak generally within the range of eight animal- and plant-based comparison products, except for compressive stiffness, resilience, and phase angle, where the fungi-based steak ranks lowest. In contrast, our complementary sensory survey positions lion’s mane steak as the highest ranked product in seven of the twelve categories, most notably in fibrousness, fattiness, moistness, and meatiness. Kendall’s rank correlations reveal that mechanical tests can predict certain sensory attributes such as softness and brittleness–repeatably, objectively, and unbiased. Taken together, lion’s mane mushroom steak is a compelling, nutritious, and sustainable alternative to plant-based and animal meats.

## Data Availability

Data and analysis scripts are freely available at https://github.com/LivingMatterLab/CANN.

## Acknowledgments

This research was supported by the NSF Graduate Research Fellowship, by the Stanford DARE Fellowship, and by the Stanford Plant-Based Diet Initiative seed grant to Skyler St. Pierre, by the Research Foundation Flanders FWO through the doctoral fellowship SB1SE2123N to Thibault Vervenne, by the Stanford Bio-X Graduate Student Fellowship to Ethan Darwin, by the Stanford Mechanical Engineering Summer Undergraduate Research Fellowship to Lucas Boyle and Manuel Palomares, by the Stanford Sapp Family CS Bio-X Undergraduate Summer Research Fellowship to Marie Goodson and Nancy Zhang, by seed funding from Food System Innovations, by the Stanford Doerr School of Sustainability Accelerator, by the Stanford Bio-X Snack Grant, by the NSF CMMI grant 2320933, and by the ERC Advanced Grant 101141626 to Ellen Kuhl.

## Notes

### Competing Interest Statement

The authors have declared no competing interest.

### Summary of Updates

- expanded statistical analysis in Section 3.4, "Sensory Analysis" - added Section 3.6, "Principal component analysis" - added Figure 9, "Principal component analysis" - adjusted Discussion to include updated results

